# Arp2/3 Complex Activity Enables Nuclear YAP for Naïve Pluripotency of Human Embryonic Stem Cells

**DOI:** 10.1101/2023.01.16.524285

**Authors:** Nathaniel P. Meyer, Tania Singh, Matthew L. Kutys, Todd Nystul, Diane L. Barber

## Abstract

Our understanding of transitions of human embryonic stem cells between distinct stages of pluripotency relies predominantly on regulation by transcriptional and epigenetic programs with limited insight on the role of established morphological changes. We report remodeling of the actin cytoskeleton of human embryonic stem cells (hESCs) as they transition from primed to naïve pluripotency that includes assembly of a ring of contractile actin filaments encapsulating colonies of naïve hESCs. Activity of the Arp2/3 complex is required for the actin ring, uniform cell mechanics within naïve colonies, nuclear translocation of the Hippo pathway effectors YAP and TAZ, and effective transition to naïve pluripotency. RNA-sequencing analysis confirms that Arp2/3 complex activity regulates Hippo signaling in hESCs, and impaired naïve pluripotency with inhibited Arp2/3 complex activity is rescued by expressing a constitutively active, nuclear-localized YAP-S127A. These new findings on the cell biology of hESCs reveal a mechanism for cytoskeletal dynamics coordinating cell mechanics to regulate gene expression and facilitate transitions between pluripotency states.

## Introduction

Derivation of clonal pluripotent stem cells (PSCs) from embryos yields cells with a spectrum of pluripotent states, dependent on developmental progression of the embryo, species, and culture condition. Clonal mouse embryonic stem cells (mESCs) represent a ground-state of pluripotency and closely recapitulate the naïve blastocyst from which they are isolated^1^. In contrast, clonal human and other primate PSCs, as conventionally isolated and maintained, exist in a primed state of pluripotency and more closely resemble the post-implantation epiblast^2^. To study the naïve state of clonal human PSCs, culture conditions were developed that dedifferentiate primed human embryonic stem cells (hESCs) to a naïve state of pluripotency^3–6^. Development of culture conditions that convert and sustain a naïve pluripotent state in human PSCs has provided an opportunity to study human development before gastrulation^7^.

These *in vitro* models of naïve pluripotency have provided insights on the transcriptomic, epigenetic, and proteomic programs that maintain a functional naïve pluripotency state in stem cells^4, 8, 9^. We have limited understanding, however, of how established morphological changes during the transit from primed to naive states are regulated and whether morphological changes regulate state transitions. It is the cytoskeleton, an intracellular network of proteins, which responds to the external microenvironment and facilitates changes in cell behavior such as cell shape. This network is primarily composed of microtubules, intermediate filaments, and actin filaments. We know that pluripotent stem cell fate is intricately regulated by biophysical cues primarily transmitted through the actin cytoskeleton, which control gene expression, proliferation, and differentiation^10^. The corresponding coordinated changes in cell shape are essential for developmental embryogenesis and are largely mediated by the actin cytoskeleton^11^. Accordingly, mechanoregulation has been studied for roles in exit from the pluripotent state toward targeted cell fates including endodermal^12^, ectodermal^13^, and mesodermal^14^ lineages^15^. Directly targeting actin filament dynamics has also been shown to regulate pluripotent stem cell fate ^16–18^. Other components of the cytoskeleton such as microtubules and intermediate filaments have also been established to modulate stem cell behavior although studies have primarily focused recently on how they impact nucleus morphology and activity^19–21^.

Actin-associated proteins, including β-catenin for enabling Wnt pathway activity and ezrin-radixin-moesin (ERM) proteins for enabling tensional forces at the plasma membrane, facilitate maintenance of the naïve pluripotent state in mESCs^22^. During murine preimplantation development, actin filaments generate mechanical forces that contribute to differentiation throughout the blastocyst stage by modulating mechanosensitive signaling pathways such as Hippo signaling^23, 24^. These actin structures allow cells within the developing blastocyst to organize based on contractility, coupling mechanosensing and fate specification^25^. Despite evidence that morphological changes and actin filament remodeling determine naïve pluripotency during mouse development, their roles in hESC naïve pluripotency remain unclear.

In asking the role of morphological changes during hESC dedifferentiation to a naïve state of pluripotency, we identified the assembly of a ring of contractile actin filaments encapsulating naïve but not primed colonies that is tethered to adherens junctions and decorated with phosphorylated myosin light chain (pMLC) and moesin. We found that nucleating activity of the Arp2/3 complex but not formins is necessary for the actin ring, naïve cell mechanics, including decreased cell-substrate tensional forces and colony formation, and transition to naïve pluripotency. RNAseq analysis revealed a role for Hippo pathway signaling in Arp2/3 regulated naïve pluripotency, which we confirmed by showing increased nuclear localization of the transcriptional co-activators YAP and TAZ in naïve compared with primed hESCs that is blocked by inhibiting Arp2/3 complex activity. Consistent with these findings, naïve pluripotency that is blocked with inhibiting Arp2/3 complex activity is restored by expressing a nuclear-localized non-phosphorylatable YAP (YAP-S127A)^26^. These data provide new mechanistic insights on how actin filament dynamics regulates the naïve state of hESCs pluripotency and the integration between actin filament remodeling and pluripotency.

## Results

### Actin Filament Remodeling as hESCs Transition to a Naïve State

For morphological analysis of pluripotency states, HUES8 primed human embryonic stem cells (hESCs) were grown on Matrigel and dedifferentiated to naïve pluripotency using previously reported conditions in an mTeSR-based medium supplemented with ERK (PD0325901) and GSK3 (CHIR99021) inhibitors, the adenylyl cyclase activator forskolin, human leukemia inhibitory factor (LIF), basic fibroblast growth factor (bFGF), and ascorbic acid ^4, 27^. We confirmed transition to a naïve state by showing that colonies have prominent doming by day 6 of dedifferentiation and increased expression of pluripotency markers DNMT3L, DPPA3, KLF2, and KLF4, as determined by rt-PCR (Fig. 1A-B). Staining fixed cells for actin filaments with phalloidin revealed that naïve but not primed colonies had a ring of bundled actin filaments at the colony periphery (Fig. 1C). These supracellular actin rings also formed around colonies of naïve H9 cells and naïve WTC11 iPSCs (Supp. Fig. 1A) as well as HUES8 cells dedifferentiated by alternative medium supplements (Supp. Fig. 1B). Moreover, the actin ring assembled independently of naïve colony size (Supp. Fig. 1C).

**Figure 1.**
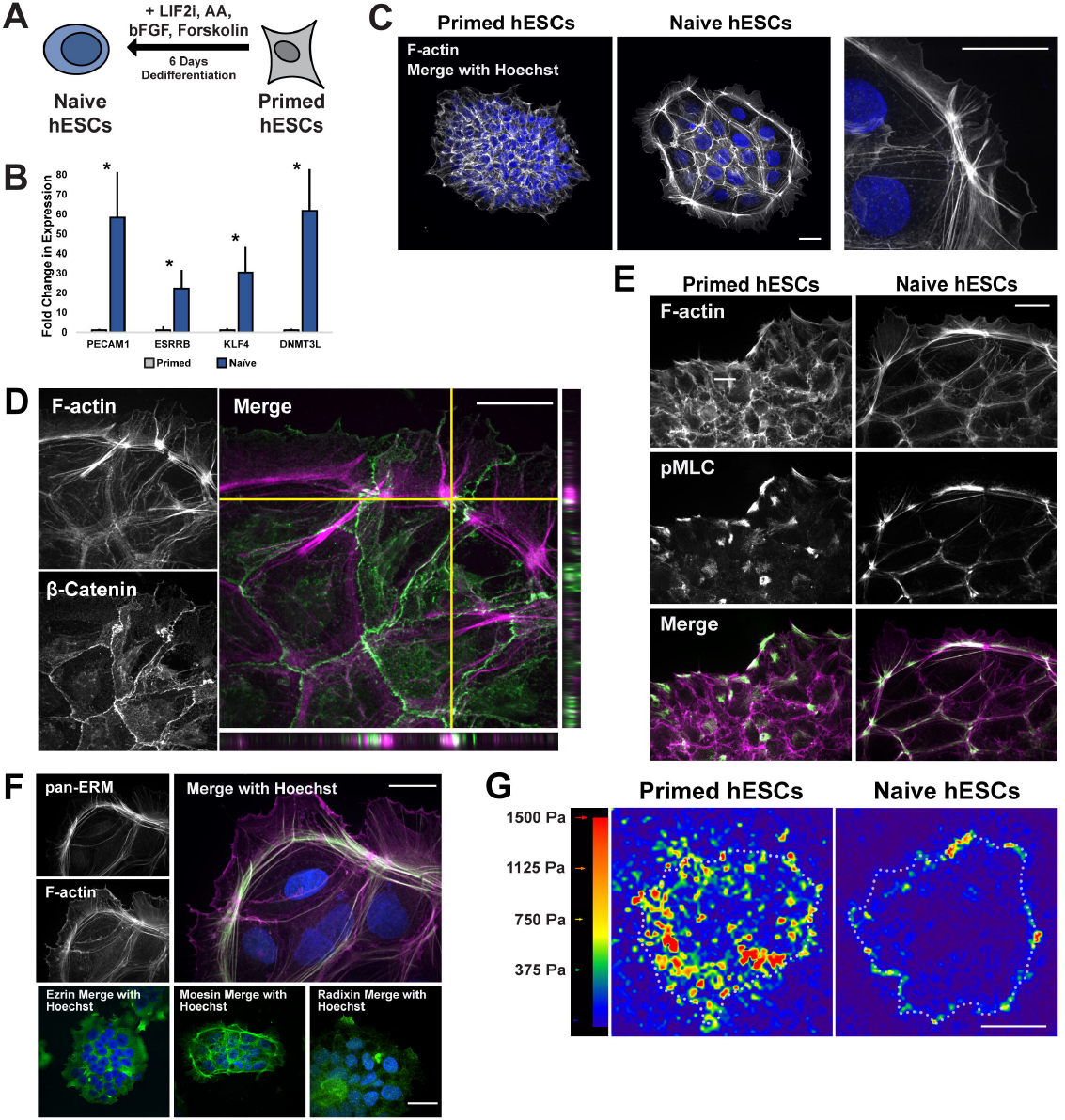
Dedifferentiation of primed hESCs to naïve pluripotency includes F-actin filament remodeling and the formation of an actin ring. **A**, Schematic of the dedifferentiation process from primed to naïve human embryonic stem cells (hESCs). **B**, Confirmation of dedifferentiation indicated by increased expression of pluripotency genes associated with a naïve state as determined by qPCR. Data represent the means ± S.E.M. normalized to Oct4 (n=3 separate cell preparations). *P* values were calculated using a two-tailed Student’s *t*-test. **C-G** Images of primed and naïve stem cells stained or immunolabeled for actin cytoskeleton components. **C**, Confocal (left and middle) and super-resolution (right) images of hESCs stained for F-actin with phalloidin (white) and Hoechst (blue) show a bundled actin filament ring encapsulating colonies of naïve but not primed cells. **D-F**, Confocal images of naïve hESCs immunolabeled for β-catenin (**D**), pMLC (**E**), and ERM proteins (**F**) and stained for F-actin with phalloidin (magenta) demonstrating cytoskeletal remodeling. Scale bars, 25 µM. **G**, Representative stress maps generated by traction force microscopy (TFM). Dotted outlines indicate colony boarders. Scale bar, 50 µM.

A similar actin ring is reported to encircle colonies of clonal human pluripotent stem cells to provide a mechanosensitive element linked to focal adhesions^28^. The actin filament ring we observed around naïve hESC colonies was instead tethered to adherens junctions, as indicated by co-labeling for β-catenin, with separated interdigitated adherens junctions suggesting a contractile force (Fig. 1D, crosshairs). The contractile property of the ring was also suggested by the actin ring being decorated with phosphorylated myosin light chain (pMLC) as determined by immunolabeling (Fig. 1E). In contrast, primed hESC colonies had irregular aggregates of pMLC with limited co-localization with actin filaments. Immunolabeling with pan-ERM antibodies showed co-localization with the actin filament ring in naïve cells, and ERM-specific antibodies revealed co-localization of moesin but not ezrin or radixin (Fig. 1F, Supp. Fig. 1D). Together, these data indicate a contractile actin ring surrounding naïve but not primed hESC colonies.

The supracellular nature of the actin ring and differential pMLC labeling between naïve but not primed hESC colonies suggested a potential difference in colony mechanics, which we determined by using traction force microscopy. Increased cell-matrix traction forces are associated with destabilized adherens junctions in epithelial monolayers^29, 30^. Consistent with pMLC localization, primed colonies exhibited elevated cell-substrate tractions that were distributed throughout the colony (Fig. 1G, left; Supp. Fig. 1E). In contrast, naïve colonies exhibited overall low magnitude cell-substrate tractions that were localized to the colony periphery and largely absent from the colony interior (Fig. 1G, right; Supp. Fig. 1E), suggesting decreased cell-substrate tensional force and a likely shift to more stabilized cell-cell forces. Along with pMLC localization, these low tractions are consistent with uniform cell-cell adhesion in naïve hESC colonies. Together these data identify a significant reorganization of the actin cytoskeleton during transition to a naïve state of pluripotency that includes the assembly of a contractile actin ring surrounding naïve cell colonies, coincident with attenuated cell-substrate traction forces and a transition to enhanced cell-cell junction traction force within the colony unit.

### Arp2/3 Complex Activity is Necessary for Transition of hESC to Naïve Pluripotency

The assembly of an actin ring in naïve but not primed human embryonic stem cell (hESC) colonies led us to ask whether the actin ring has a functional significance in the transition to naïve pluripotency. New actin filaments are predominantly generated by two distinct nucleators, the Arp2/3 complex, which generates branched filaments, and formins, which generate unbranched filaments^31^. We found that the actin ring assembled when naïve cells are generated in the presence of SMIFH2, a broad-spectrum inhibitor of formin activity^32, 33^ but not CK666, a selective pharmacological inhibitor of Arp2/3 complex activity^34, 35^ (Fig. 2A). Additionally, CK666 blocked increased expression of markers of naïve pluripotency seen in controls, determined by qPCR of PECAM1, ESRRB, KLF4, and DNMT3L (Fig. 2B). To eliminate the possibility that CK666 led cells to exit pluripotency and differentiate, we immunolabeled for the general pluripotency markers OCT4 and SOX2 and found that CK666 treated cells remained broadly pluripotent (Supp. Fig. 2). To further assess the pluripotent state of cells dedifferentiated in the presence of CK666, we immunolabeled for the primed pluripotent marker SSEA3^36^. In controls, SSEA3 expression significantly decreased with dedifferentiation, as previously reported^37^ but not with CK666 (Fig. 2C, D). Additionally, the naïve pluripotency marker KLF4^38^ translocated from the cytoplasm to the nucleus with control dedifferentiation but not in the presence of CK666 (Fig. 2C, E).

**Figure 2.**
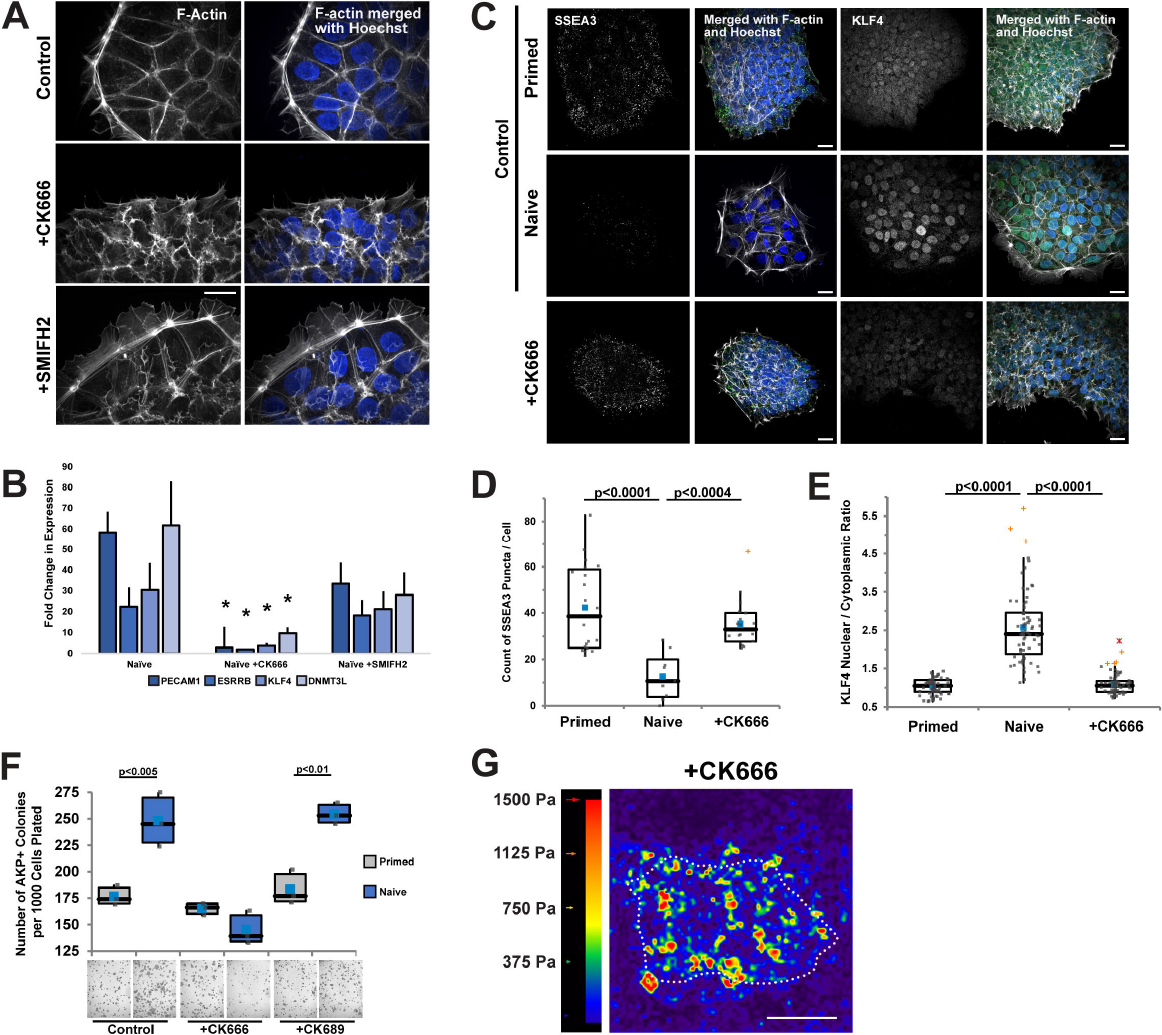
Inhibiting Arp2/3 complex but not formin activity blocks formation of an actin ring and dedifferentiation to naïve pluripotency. **A**, Confocal images of D6 naïve hESCs maintained in the absence (control) or presence of 80 µM CK666 or 50 µM SMIFH2 and stained for F-actin with phalloidin and nuclei with Hoechst. **B**, Expression of the indicated pluripotency transcripts determined by qPCR at D6 of dedifferentiation in the absence (Control) or presence of CK666 or SMIFH2. The Arp2/3-complex activity inhibitor CK666 impairs upregulation of pluripotency genes used to identify naïve pluripotency. Data are the means ± S.E.M. of 3 determinations normalized to Oct4. *P* values were calculated using a two-tailed Student’s *t*-test. **C-E**, Confocal images of control primed and naïve hESCs and D6 cells dedifferentiated in the presence of CK666 immunolabeled for the primed marker SSEA3, quantified in (**D**) and the naïve marker KLF4, quantified in (**E**). Box plots in (**D**) and (**E**) show median, first and third quartile, with whiskers extending to observations within 1.5 times the interquartile range. **F**, Clonogenicity, determined by alkaline phosphatase positive colonies (quantified in top panel and representative brightfield images in bottom panel) in control primed and naïve hESC as well as dedifferentiated in the presence of CK666 or 80 µM CK689, an inactive analog of CK666. Data are the means ± S.E.M. normalized to the number of cells plated in 3 separate determinations. Box plots are as described in (**D,E**) with *P* values calculated using a two-tailed Student’s *t*-test. Scale bars, 25 µM. **G**, Representative stress maps generated by traction force microscopy (TFM). Dotted outlines indicate colony boarders. Scale bar, 50 µM.

We further tested for a functional naïve pluripotent state by staining for alkaline phosphatase (AKP) and scoring for colony formation, which indicates the capacity for clonogenic expansion and self-renewal^39^. Primed and naïve hESCs were passaged and plated at clonogenic cell numbers and maintained for five days without or with CK666. In controls, colony formation was greater in naïve compared with primed hESC, as previously reported^40^. However, with CK666, but not CK689, an inactive analog of CK666, there was no increase in colony formation in naïve compared with primed cells (Fig. 2F). Additionally, traction force microscopy revealed that elevated cell-substrate tractions throughout colonies of primed but not naïve cells (Fig. 1G), were retained when hESCs were dedifferentiated in the presence of CK666 (Fig. 2G; Supp. Fig. 2B). These data identify an essential role for the Arp2/3 complex in promoting an actin filament ring and uniform naïve colony mechanics as well as acquiring a naïve pluripotent state in hESCs.

### Arp2/3 Complex Activity Enables Active YAP for Naïve Pluripotency

To understand how Arp2/3 complex activity affects the transcriptional circuitry required for naïve pluripotency, we performed bulk RNA-sequencing (RNAseq) on primed cells, naïve cells, and primed human embryonic stem cells (hESCs) dedifferentiated in the presence of CK666 (Fig. 3A). We found that primed, naïve, and CK666-treated cells had a total of 12,817 differentially expressed genes (DEGs) with an adjusted *pval <0.05*. Of these DEGs, 182 were unique to control primed cells compared with control naïve cells and were not differentially expressed in CK666-treated cells; CK666-treated cells compared with control primed or control naïve cells had 102 and 502 DEGs, respectively. To determine the transcriptional networks involved in the dedifferentiation from primed to naïve pluripotency, we identified KEGG pathways in control naïve dedifferentiation that revealed Hippo signaling as the top candidate (Fig. 3B). Additionally, transcription factor binding motif analysis identified TEAD2 as a top candidate, which is a downstream effector of Hippo signaling (Fig. 3C).

**Figure 3.**
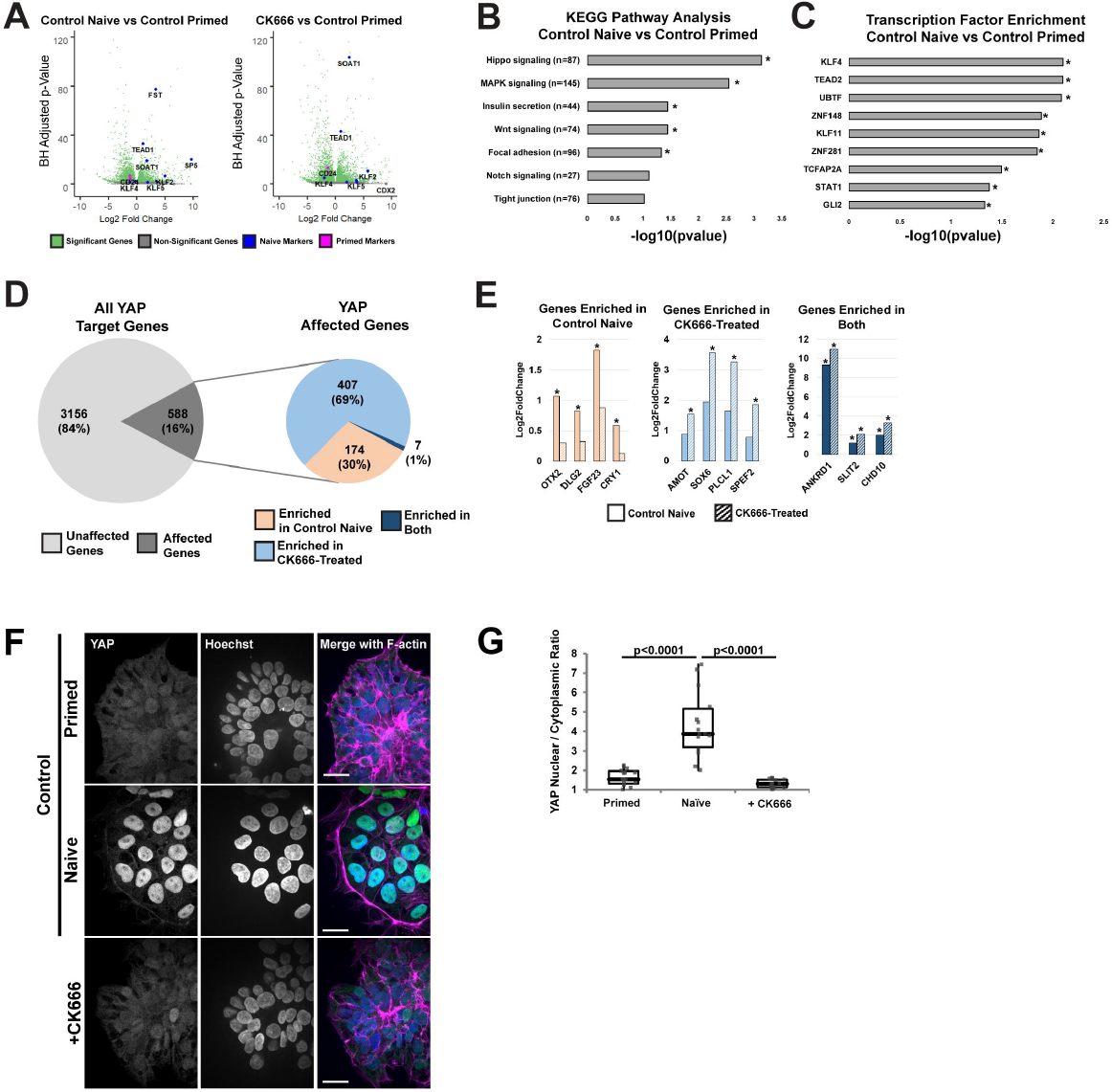
Inhibiting Arp2/3 complex activity disrupts Hippo signaling in naïve hESCs. **A**, Volcano plots showing transcriptome fold-changes (*padj)* of dedifferentiations in the absence (Control Naïve) or presence of CK666-treated dedifferentiations compared with primed hESCs. Each dot represents a single gene, with significant genes (padj &lt; 0.05) in green). Notable primed and naïve markers are depicted in red and blue, respectively. **B**,**C** KEGG pathway analysis (**B**) and transcription factor enrichment analysis (**C**) of control primed and naïve hESCs. The number of DEGs indicated in each pathway is displayed and asterisks indicate significantly enriched pathways (*p* &lt; 0.05). **D**, Unbiased screening of all known YAP target genes in dedifferentiated cells in the absence (Control Naive) and presence of CK666. Affected genes were further analyzed to indicate whether they are enriched in control, CK666-treated, or both conditions when compared with primed controls. **E**, Expression of selected YAP target genes from bulk RNAseq, with asterisks indicating significant difference (*padj* &lt; 0.05). **F**, Representative confocal images of control primed and naïve hESCs and naïve hESCs generated in the presence of 80 µM CK666 immunolabeled for YAP and stained for nuclei with Hoechst and F-actin with phalloidin. **G**, Quantification of nuclear to cytoplasmic ratio of YAP from images as shown in (**F**). Box plots show median, first and third quartile, with whiskers extending to observations within 1.5 times the interquartile range. Data are from 5 separate cell preparations with *P* values calculated using a two-tailed Student’s *t*-test. Scale bars, 25 µM.

The Hippo effector protein YAP is a known regulator of the human naïve pluripotent state, with overexpression of YAP in pluripotent stem cells promoting the acquisition of naïve pluripotency^27^. Although actin filament dynamics, including a contractile ring of actin, regulate YAP signaling ^41, 42^, to our knowledge a role for Arp2/3 complex activity regulating YAP or TAZ activity in human naïve pluripotency has not been reported. For an unbiased global analysis of known YAP target genes, we used two publicly available datasets^43, 44^ and found that of the 3,744 YAP target genes identified in our RNAseq dataset, 3156 (84%) were not differentially expressed in any condition and 588 (16%) were enriched in one or multiple conditions. Of those 588 enriched YAP-target genes, 174 (30%) were significantly enriched in the control naïve dedifferentiation condition versus the control primed condition; 407 (69%) were significantly enriched among CK666-treated dedifferentiation condition versus the control primed condition; and 7 (1%) were significantly enriched in both conditions versus the control primed condition (Fig. 3D, adjusted pval >0.05).

Of the genes significantly enriched in the control naïve condition compared with the control primed condition, known naïve pluripotency markers such as OTX2, DLG2, and CRY1 were significantly upregulated; these naïve markers failed to significantly increase in the CK666-treated condition (Fig 3E, left). As expected, genes significantly enriched among both DEG lists include known YAP and Hippo targets such as ANKRD1, SLIT2, and CHD10. (Fig. 3E, right). Genes significantly enriched among the CK666-treated condition include the negative Hippo regulator AMOT^45^, and lineage-commitment genes such as SOX6 and SPEF2 (Fig. 3E, middle). These data suggested a Hippo signaling pathway program, driven by mediators such as YAP, occurs during dedifferentiation to naïve pluripotency but is disrupted by inhibiting Arp2/3-complex activity. To verify this prediction, we immunolabeled cells to determine YAP localization and found increased nuclear to cytoplasmic ratios of YAP (Fig. 3F-G) and TAZ (Supp. Fig. 3) with control dedifferentiation that was blocked by CK666.

Consistent with these data, during preimplantation development, actin filaments and associated proteins generate mechanical forces that contribute to differentiation throughout the blastocyst stage through modulation of mechanosensitive pathways such as Hippo signaling^23, 24^. These actin structures allow cells within the developing blastocyst to organize based on contractility, coupling mechanosensing and fate specification^25^. Therefore, we hypothesized that Arp2/3 complex activity facilitated naïve dedifferentiation through increasing YAP nuclear localization. To test this prediction, we asked whether primed hESCs stably expressing a constitutively active, nuclear-localized form of YAP (YAP-S127A) could restore naïve dedifferentiation in the presence of CK666. Accordingly, we observed that two immunofluorescence-based markers of the naïve state, increased nuclear localization of KLF4 (Fig. 4A-B) and decreased SSEA3 (Fig. 4C-D), were rescued by overexpression YAP-S127A in the presence of CK666. In contrast, acquisition of a naïve state of pluripotency remained blocked with CK666 treatment in cells overexpressing wildtype YAP (WT YAP) (Fig. 4A-D). Additionally, YAP-S127A rescued colony formation, a functional form of naïve pluripotency, that was inhibited with CK666 (Fig. 4E).

**Figure 4.**
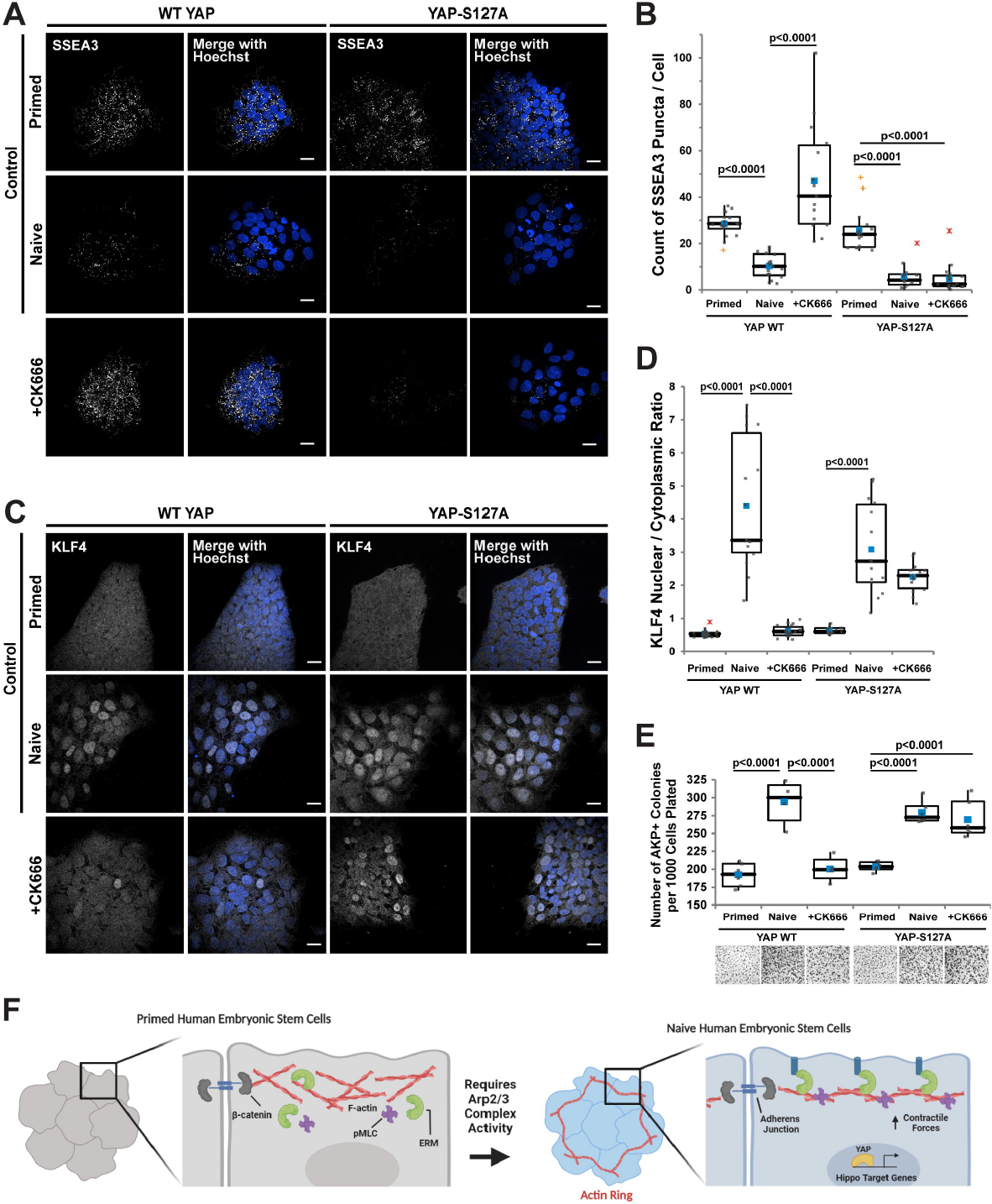
Overexpression of YAP-S127A rescues naïve pluripotency blocked with inhibiting Arp2/3 complex activity. **A, C**, Representative confocal images of control primed, control naïve cells, and cells dedifferentiated in the presence of CK666 with or without stably overexpressing YAP WT or YAP-S127A immunolabeled for the primed marker SSEA3 (**A**) or the naïve marker KLF4 (**C**) and stained for nuclei with Hoechst and F-actin with phalloidin. **B, D**, Images as in **A** and **C** were used to quantify, respectively, the number of SSEA3 puncta (**B**) and the nuclear to cytoplasmic ratio of KLF4 (**D**). Box are plots as described for Figure 2. **E**, Clonogenicity, determined by alkaline phosphatase positive colonies (quantified in top panel and representative brightfield images in bottom panel) in control primed and naïve hESC, and dedifferentiated in the presence of CK666 with stably expressed YAP WT or YAP-S127A. Data are the means ± S.E.M. normalized to the number of cells plated from 3 separate determinations, with box plots as described for Figure 2 d, e and *P* values are calculated using a two-tailed Student’s *t*-test. Scale bars, 25 µM. **F**, Model illustrating cytoskeletal remodeling that occurs during dedifferentiation to a naïve state of pluripotency. Successful dedifferentiation to naïve pluripotency includes the formation of an actin ring structure, uniformity of colony mechanics, recruitment of actin-binding proteins known to play roles in pluripotency, and the regulation of Hippo signaling.

## Discussion

Our new findings support a model in which naïve pluripotency is characterized by an Arp2/3 complex-dependent remodeling of the actin cytoskeleton that includes formation of a contractile supracellular actin ring enclosing naïve colonies and establishment of uniformity in colony mechanics likely enabled by the actin ring being physically associated with β-catenin and moesin, which are known to play roles in pluripotency^46, 47^ (Fig. 4F). Moreover, Arp2/3 activity facilitates dedifferentiation to a naïve state of pluripotency through promoting nuclear translocation of YAP and regulating Hippo target gene expression. Consistent with these findings, naïve pluripotency that is blocked with inhibited Arp2/3 complex activity is restored by expressing a constitutively active, nuclear-localized YAP-S127A.

Our findings are distinct from those on a contractile actin filament ring that assembles around colonies of primed pluripotent stem cells^28^ and Xenopus neural crest cells^48^, which functions to enhance cell-substrate adhesion and migratory capacity, respectively. Our findings also highlight distinct differences between murine and human embryonic stem cells. Cells within the ICM of mouse blastocysts exclude YAP from the nucleus whereas cells within the ICM of human blastocysts maintain nuclear YAP^27, 49^. This difference in YAP localization is retained *in vitro*, with murine naïve pluripotent stem cells (PSCs) having predominantly cytosolic YAP ^50^, and human naïve PSCs having predominantly nuclear YAP (Fig. 3F-G). How this difference in YAP localization occurs between mouse and human is unknown, although a number of cytoskeletal factors regulate YAP localization, including stability of the actin cytoskeleton, contractility, and mechanical regulators such as ERM proteins^42^. The role of the actin cytoskeleton in the exit from the pluripotent state has also highlighted how actin dynamics may facilitate cell fate decisions. For example, cells located at the colony edge of primed human embryonic stem cells (hESCs) have distinct cytoskeletal dynamics and are uniquely poised to exit pluripotency and differentiate^51, 52^. Positional differences in differentiation potential such as these have been proposed as a mechanism executed in early embryo symmetry breaking with rearrangement of the actin cytoskeleton being required for the first cell fate decision in the blastocyst^53, 54^. Thus, it may be possible that the contractile actin ring we observe at the edge of naïve colonies (Fig 1. C) functions as a hub for regulating cell fate dynamics through similar mechanisms as first cell fate decision including modulation of mechanosensitive signaling such as YAP and through pathways such as Hippo.

Further highlighting differences between human and mouse embryonic stem cells, we recently reported that Arp2/3 complex activity is necessary for the differentiation of clonal mouse naïve PSCs to the primed epiblast state, which is in part mediated by translocation of myocardin-related transcription factor MRTF from the cytosol to the nucleus^55^. Additionally, a recent report suggests that Arp2/3 complex activity may form a positive feedback loop with YAP-TEAD1 transcriptional activity controlling cytoskeletal reorganization ^56^; thus Arp2/3 complex activity may regulate naïve pluripotency at multiple stages including initially to reorganize the actin cytoskeleton, but also during maintenance of naïve pluripotency through regulating YAP localization and hence activity.

Our findings increase our understanding of actin dynamics and cell mechanics as regulators of cell fate transitions. A role for contractile actin filaments as a mechanoresponsive element for pluripotency states is well established^57^, and our work identifies cytoskeletal dynamics essential for uniform colony mechanics and the naïve pluripotent state, the role of Arp2/3 complex activity, and YAP/TAZ activity as a promising target for reprogramming of hESCs and for regenerative medicine.

## METHODS

### Cell Culture

Primed human embryonic stem cell lines HUES8, H9, and WTC11 were maintained on Matrigel (Corning Life Science #354277) in feeder-free mTeSR-1 medium (STEMCELL Technologies # 85850) at 37°C with 5% CO_2_ with daily medium changes. Cells were passaged approximately every 3 days, by dissociating with Accutase (STEMCELL Technologies #07920) and including the Rho-associated coiled-coil kinase (ROCKi) inhibitor Y-276932 (10 µM; Selleckchem #S1049) in the plating medium to facilitate survival. All cell lines were routinely confirmed to be negative for mycoplasma by testing with a MycoAlert Mycoplasma Detection Kit (Lonza # LT07-701).

### Generation of Naïve hESCs

Dedifferentiation was completed using previously published methods^4, 27^. In brief, cells were plated at a density of 10,000 cells per cm^2^ in the presence of ROCKi (10µM). After 24 h, cells were washed 3 times with PBS and incubated in naïve dedifferentiation medium of mTeSR-1 supplemented with 12 ng/mL bFGF (Peprotech #AF-100-18B), 1 µM PD0325901 (MEKi, Selleckchem #S1036), 3 µM CHIR99021 (GSK3βi, Selleckchem # S2924), 10 µM Forskolin (StemCell Technologies #72112), 50 ng/mL ascorbic acid (Sigma # A92902), and 1000 U recombinant human LIF (StemCell Technologies #78055). Medium was replaced daily, and cells were passaged every 3 days with Accutase. Where indicated, the naïve dedifferentiation medium 2iFL was used, which was mTeSR-1 supplemented with 0.5 μM PD0325901, 3 μM CHIR9902, 10 μM Forskolin, and 1000 U recombinant human LIF. For actin nucleator experiments, naïve dedifferentiation media was supplemented with either 80 μM CK666 (EMD Millipore #182515), 80 μM CK689 (EMD Millipore #182517), or 50 μM SMIFH2 (Sigma #S4826) throughout the entire dedifferentiation process. When passaging, media was supplemented with both ROCKi (10µM) and the appropriate inhibitor.

### qPCR

Total RNA was isolated using RNAeasy Mini Plus (Qiagen #74134) kits and cDNA was generated using iScript cDNA Synthesis kits (Bio-Rad #1708890) as per the manufacturer’s specifications. Quantitative PCR was performed using iQ SYBR Green Supermix (Bio-Rad #1708882) and analyzed on a QuantStudio six Flex Real-Time PCR System (Applied Biosystems).

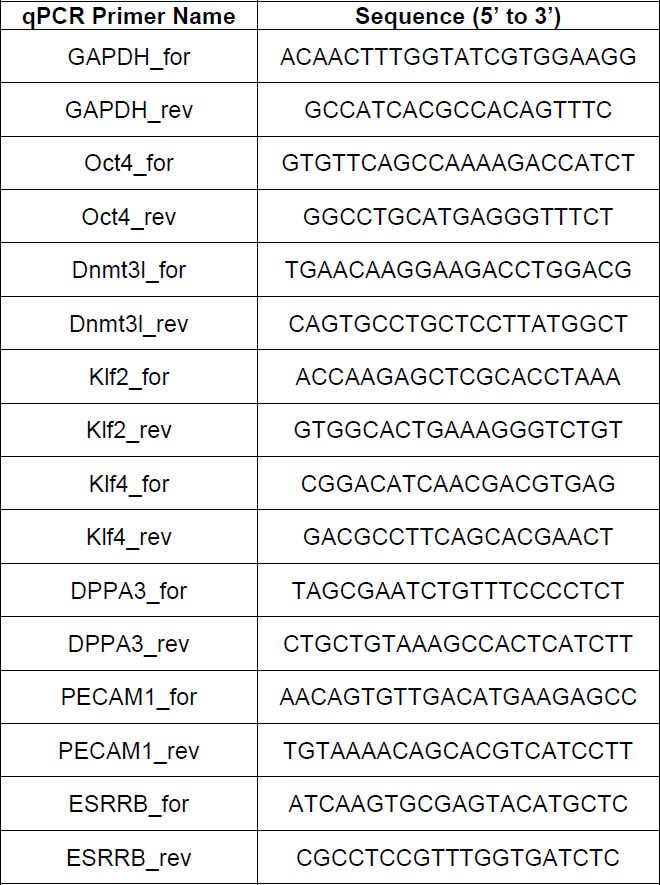

### Staining and Immunolabeling

For microscopy, cells were plated on Matrigel-coated glass coverslips prepared using an ultrasonic cleaning bath (Branson). In brief, coverslips were sonicated for 20 minutes in the presence of diluted Versa-Clear (FisherScientific #18-200-700) in double distilled H_2_O (ddH_2_O), washed three times using ddH_2_O, sonicated for 20 minutes in ddH_2_O, washed three times using ddH_2_O, and sterilized and stored in 70% ethanol. Cells were maintained for indicated times, typically 3 days, washed briefly with PBS, fixed with 4% PFA for 12 minutes at room temperature, permeabilized with 0.1% Triton X-100 in PBS for 5 minutes, and incubated with blocking buffer consisting of 0.1% Triton X-100 in PBS and 1% BSA for 1 h. Cells were then incubated with primary antibodies diluted in blocking buffer overnight at 4°C, washed with PBS three times, and incubated for 1 hour at room temperature with secondary antibodies, followed by a final PBS 3X wash, with the second wash containing Hoechst 33342 (1:10,000; Molecular Probes #H-3570) to stain nuclei. To stain for actin filaments, either rhodamine phalloidin (1:400, Invitrogen # R415) or Phalloidin-iFluor 647 (1:1000, Abcam # ab176759) was added to the secondary antibody incubation. Secondary antibodies used were Alexa Fluor 488 or 594 for the appropriate species primaries (1:500, Invitrogen #A-11037).

| **Antibody** | **Source** | **Catalog Number** | **Dilution** |
| --- | --- | --- | --- |
| β-catenin | BD Transduction | #610154 | 1:200 ICC |
| pMLC (Thr18/Ser19) | Cell Signaling | #3674 | 1:200 ICC |
| pan-ERM | Cell Signaling | #3142 | 1:400 ICC |
| Moesin | Cell Signaling | #3146 | 1:400 ICC |
| Ezrin | Cell Signaling | #3145 | 1:400 ICC |
| SSEA3 | Santa Cruz | sc-21703 | 1:200 ICC |
| KLF4 | Cell Signaling | #4038 | 1:200 ICC |
| YAP | Sigma Aldrich | HPA038885 | 1:400 ICC |
| TAZ | Sigma Aldrich | HPA039557 | 1:400 ICC |
| Oct3/4 | Santa Cruz | sc-5279 | 1:400 ICC |
| Sox2 | Cell Signaling | #3579 | 1:400 ICC |

### Confocal and Super-resolution Image Acquisition and Quantification

Cells were imaged using an inverted microscope system (Nikon Eclipse TE2000 Perfect Focus System; Nikon Instruments) equipped with a spinning-disk confocal scanner unit (CSU10; Yokogawa), a 488-nm solid-state laser (LMM5; Spectral Applied Research), and a multipoint stage (MS-2000; Applied Scientific Instruments). A CoolSnap HQ2 cooled charge-coupled camera (Photometrics) was used to take images with a camera triggered electronic shutter controlled by NIS Elements Imaging Software (Nikon) and a 60X Plan Apochromat TIRF 1.45 NA oil immersion objective equipped with a Borealis (Andor) to normalize lamination. High resolution and super-resolution images were acquired using a Yokogawa CSU-W1/SoRa spinning disk confocal system (Yokogawa) and an ORCA Fusion BT sCMOS camera (Hamamatsu) using 2×2 camera binning. Nuclear-to-cytoplasmic ratios of immunolabeled proteins and number of puncta per cell were quantified using NIS Elements Imaging Software (Nikon). Briefly, the fluorescence in the nucleus (detected by Hoescht) and in the cytoplasm were manually sampled by selection of regions-of-interest. Three regions-of-interest outside of any cell were used to calculate background fluorescence and was subtracted from both nuclear and cytoplasmic fluorescence values. The ratio of fluorescence was then determined by diving the nuclear fluorescence intensity with that of the cytoplasm for a given cell. Quantification of puncta for SSEA3 was done by creating a 3D projection of full-cell z-stacks by using NIS Elements Imaging Software. Surfaces were created using the 3D thresholding tool normalized across all images and the total number of puncta was recorded. The total number of cells was then counted, as determined by the number of Hoescht positive nuclei, and the number of puncta per cell was calculated by dividing the total number of puncta by the number of cells in each field of view.

### Traction Force Microscopy

Polyacrylamide gels of 7.9 kPa stiffness were made by adjusting acrylamide and bisacrylamide stock solution (Bio-Rad Laboratories, Hercules, CA) concentrations. A solution of 40% acrylamide, 2% bisacrylamide and 1xPBS was polymerized by adding tetramethylethylene diamine (Fisher BioReagents) and 1% ammonium persulfate. A droplet of the gel solution supplemented with 0.2 μm fluorescent beads solution (Molecular Probe, Fisher Scientific) was deposited on a quartz slide (Fisher Scientific) and covered with a 25-mm glass (Fisher) coverslip pretreated with 3-aminopropyltrimethoxysilane (Sigma-Aldrich) and glutaraldehyde (Sigma-Aldrich). After polymerization, the gel surface attached to the quartz slide was functionalized with Matrigel via polydopamine. The gel was sterilized and stored in 1X PBS before cell seeding. The traction forces exerted by colonies on the polyacrylamide gel substrates were computed by measuring the displacement of fluorescent beads embedded within the gel. Briefly, images of bead motion near the substrate surface, distributed in and around the contact region of a single cell (before and after cell detachment with 10% sodium dodecyl sulfate), were acquired with Yokogawa CSU-21/Zeiss Axiovert 200M inverted spinning disk microscope with a Zeiss LD C-Apochromat 40X, 1.1 NA water-immersion objective and an Evolve EMCCD camera (Photometrics). The traction stress vector fields were generated using an open-source package of FIJI plugins (https://sites.google.com/site/qingzongtseng/tfm).

### Colony Formation Assay

To determine clonogenic potential, cells were dissociated with Accutase and plated on Matrigel-coated 6-well dishes at a density of 1,000 cells per cm^2^ in the presence of ROCKi (10 µM). Five days after plating, cells were stained for alkaline phosphatase as per the manufacturer’s protocol (StemAb Alkaline Phosphatase Staining Kit II, ReproCell #00-0055) and imaged using a Leica DFC 7000t microscope. To quantify the number of alkaline phosphatase positive colonies, images were analyzed using Fiji^58^.

### Library Preparation and RNA Sequencing

RNA was extracted using RNeasy Mini kits (Qiagen) according to the manufacturer’s instructions and concentrations were determined by NanoDrop. Library preparation and RNA sequencing were performed by Novogene Co. Ltd (USA). Briefly, RNA purity was measured using a NanoPhotometer spectrophotometer (IMPLEN). RNA integrity and quantity were determined using a Bioanalyzer 2100 system (Agilent Technologies). Three paired biological replicate libraries were prepared for each condition, with each library generated with 1 µg of RNA per sample. Sequencing libraries were generated using NEBNext Ultra RNA Library Prep Kit for Illumina (NEB) following manufacturer’s recommendations and index codes were added to attribute sequences to each sample. Briefly, mRNA was purified from total RNA using poly-T oligo-attached magnetic beads. Fragmentation was carried out using divalent cations under elevated temperature in NEBNext First Strand Synthesis Reaction Buffer (5X). First strand cDNA was synthesized using random hexamer primer and M-MuLV Reverse Transcriptase (RNase H-). Second strand cDNA synthesis was subsequently performed using DNA Polymerase I and RNase H. Remaining overhangs were converted into blunt ends via exonuclease/polymerase activities. After adenylation of 3’ ends of DNA fragments, NEBNext Adaptor with hairpin loop structure were ligated to prepare for hybridization. In order to select cDNA fragments of preferentially 150∼200 bp in length, the library fragments were purified with AMPure XP system (Beckman Coulter, Beverly, USA). Then 3 µl USER Enzyme (NEB, USA) was used with size-selected, adaptorligated cDNA at 37 °C for 15 min followed by 5 min at 95°C before PCR. Then PCR was performed with Phusion High-Fidelity DNA polymerase, Universal PCR primers and Index (X) Primer. At last, PCR products were purified (AMPure XP system) and library quality was assessed on the Agilent Bioanalyzer 2100 system.

### RNA Sequencing Analysis

Raw data (raw reads) were processed through fastp to remove adapters, poly-N sequences, and reads with low quality. Q20, Q30 and GC content of the clean data were calculated and found to be within the normal range. All the downstream analyses were based on the clean data with high quality. Reference genome (ID: 51) and gene model annotation files were downloaded from genome website browser (NCBI) directly. Paired-end clean reads were aligned to the reference genome using the Spliced Transcripts Alignment to a Reference (STAR) software. FeatureCounts was used to count the read numbers mapped of each gene. And then RPKM of each gene was calculated based on the length of the gene and reads count mapped to this gene. Differential expression analysis was performed using DESeq2 R package. The resulting P values were adjusted using the Benjamini and Hochberg’s approach for controlling the False Discovery Rate (FDR). Genes with a padj < 0.05 found by DESeq2 were assigned as differentially expressed. The R package clusterProfiler was used to test the statistical enrichment of differential expression genes in KEGG pathways. KEGG terms with padj < 0.05 were considered significant enrichment. Transcription factor binding motif analysis was performed using Enrichr^59, 60^. To investigate YAP target gene expression, supplementary tables generated as previously described (Pagliari et al., 2020 and Estarás et al. 2017, see Supplemental Methods) were used to generate YAP target gene lists^43, 44^.

### Plasmids, Site-Directed Mutagenesis, and Generation of Lentivirus

pGAMA-YAP was a gift from Miguel Ramalho-Santos (Addgene plasmid #74942). Site-directed mutagenesis was performed on the pGAMA-YAP construct to create p-GAMA-YAP-S127A using the QuikChange Lightning kit (Agilent Technologies #210513). Forward primers used for the Ser-to-Ala substitution was as follows: 5′-GTTCGAGCTCATGCCTCTCCAGC-3′ and 5’-GCTGGAGAGGCATGAGCTCGAAC-3’. The pGAMA-YAP-S127A plasmid was confirmed via DNA sequencing. To prepare lentivirus, HEK293-FT (Invitrogen # R70007) cells were grown in Dulbecco’s Modified Eagle Medium (ThermoFisher #11965118) supplemented with 10% fetal bovine serum (Peak Serum #PS-FB4), non-essential amino acids (UCSF CCF #CCFGA001), pen/strep (UCSF CCF #CCFGK003), and sodium pyruvate (UCSF CCF #CCFGE001) and maintained at 37°C with 5% CO_2_. Lentivirus was generated according to the manufacturer’s specifications by co-transfecting HEK293-FT’s with a mixture of packaging plasmids (ViraPower Lentivirus Expression System; ThermoFisher #K497500). Briefly, 5 x 10^6^ HEK293-FT’s were seeded onto a 10 cm dish containing 10 mL of complete medium without antibiotics. After 24h, cells were transfected with a mixture of 3 µg of the lentiviral plasmid containing the gene of interest and 9 µg of the ViraPower Packaging Mix using Lipofectamine^TM^ 2000 (ThermoFisher # 11668030). At 72 hours post-transfection, supernatant was collected, filtered, and concentrated using Lenti-X Concentrator (Takarabio #631231). Concentrated viral supernatant was aliquoted and stored at −80°C.

To generate hESC lines stably expressing WT-YAP and YAP-S127A, primed HUES8 cells were grown to approximately 60% confluency in one well of a 6-well plate. Medium was aspirated, washed once with PBS, and cells were then fed with 1mL fresh media containing 2 µg of polybrene (Millipore Sigma #TR-1003-G), and incubated for 15 minutes at 37°C. Concentrated virus supernatant (100 µL) was added and after 6-8 hours 1 mL of fresh medium was added. After 36 h, viral particles were removed by replacing medium. Three days after virus infection, hESCs were passaged and expanded to three wells in a 6 well-plate. After reaching ∼75% confluency, hESCs and sorted for high mCherry expression by using a BD FACS Aria3u, with sorted cell maintain with pen/strep for 3 days and then maintained in standard antibiotic-free TESR+ media.

---

**Supplemental Figure 1. A**, Confocal images of naïve H9 hESCs and WTC11 iPSCs stained for F-actin with phalloidin. **B**, Confocal images of naïve hESCs using an alternative dedifferentiation medium^27^. **C**, Confocal images of naïve hESCs at different colony sizes. **D**, Confocal images of primed hESCs immunolabeled with pan-ERM antibodies and stained for F-actin with phalloidin and nuclei with Hoechst. Scale bars, 25 µM. **E**, Phase contrast images of colonies used to show representative tractions for traction force microscopy in Figure 1G.

**Supplemental Figure 2. A**, Confocal images of naïve hESCs stained for F-actin with phalloidin (magenta) and Hoechst (blue) and immunolabeled for pluripotency markers Oct4 (green) and Sox2 (yellow). Scale bars, 25 µM. **B**, Phase contrast images of colonies used to show representative tractions for traction force microscopy in Figure 2G.

**Supplemental Figure 3. A**, Confocal images of D6 primed and naïve hESCs in the absence (Control) or presence of 80 µM CK666 immunolabled for TAZ (green) and stained for F-actin with phalloidin (magenta) and Hoechst (blue). **B**, Quantification of nuclear to cytoplasmic ratio of TAZ from images shown in **A**. Box plots show median, first and third quartile, with whiskers extending to observations within 1.5 times the interquartile range. Data are from 5 separate cell preparations with *P* values calculated using a two-tailed Student’s *t*-test. Scale bars, 25 µM.

