## Supplementary figures and images for "Arp2/3 Complex Activity Enables Nuclear YAP for Naïve Pluripotency of Human Embryonic Stem Cells"

### Supplemental Figure 1

#### Supplementary Figure 1. Meyer et al.

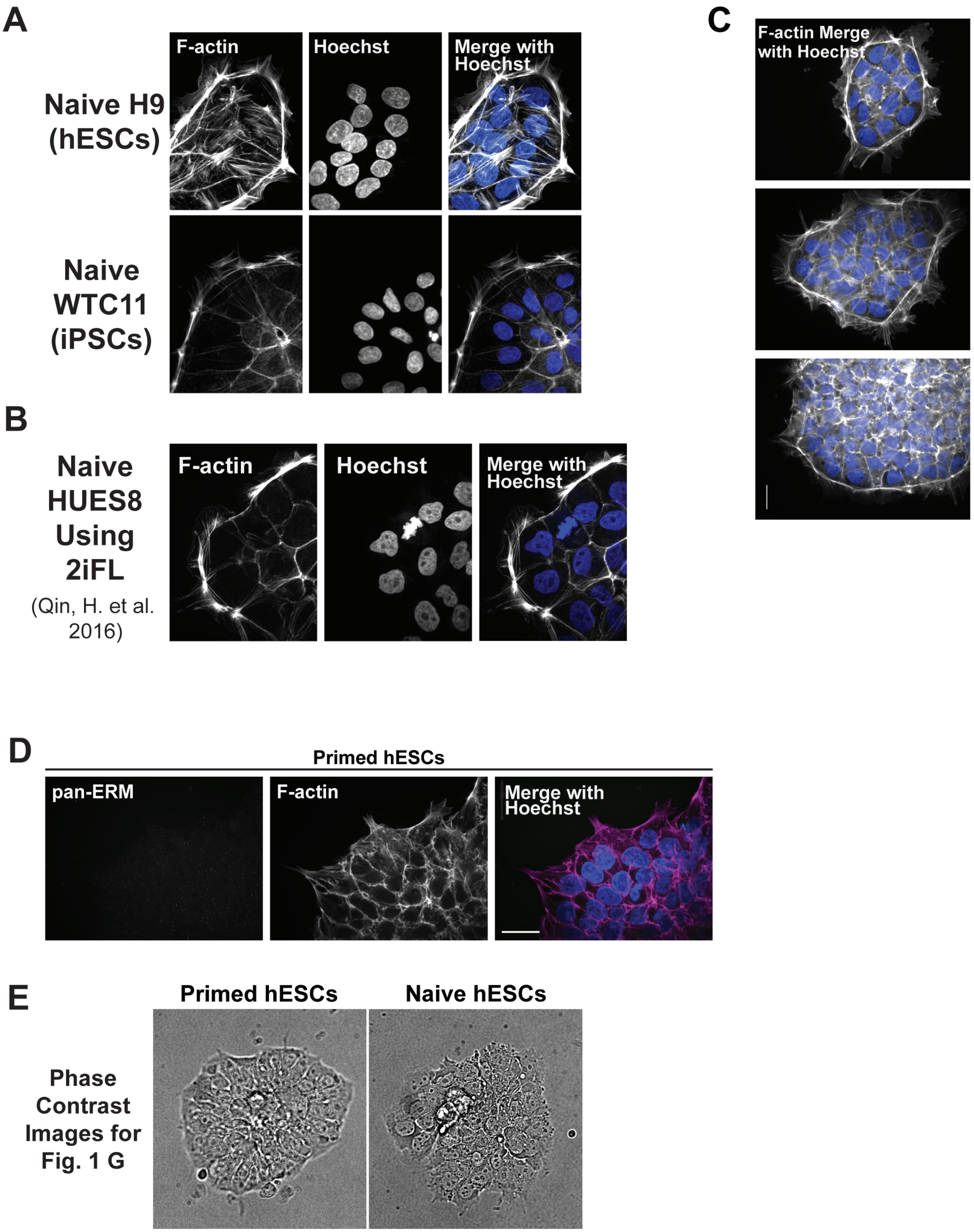

### Supplemental Figure 2

#### Supplementary Figure 2 Meyer et al.

A

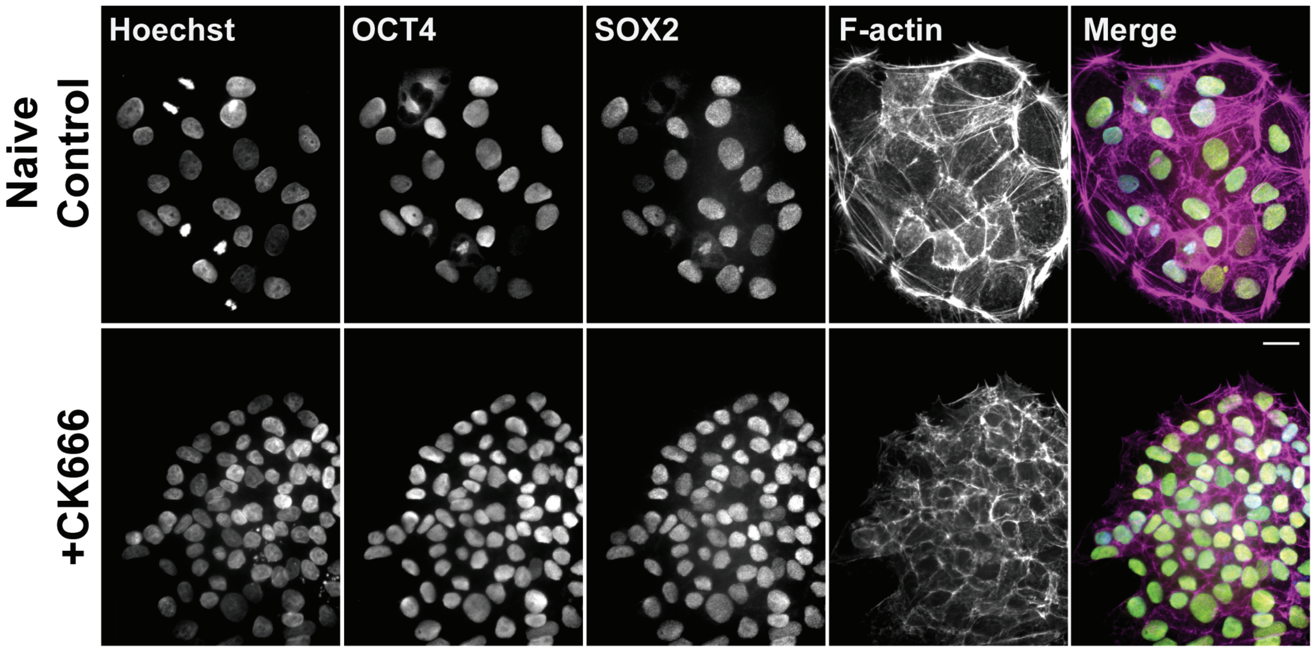


B

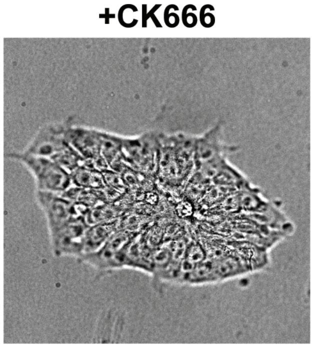


Phase Contrast Images for Fig. 2 G

### Supplemental Figure 3

#### Supplementary Figure 3. Meyer et al.

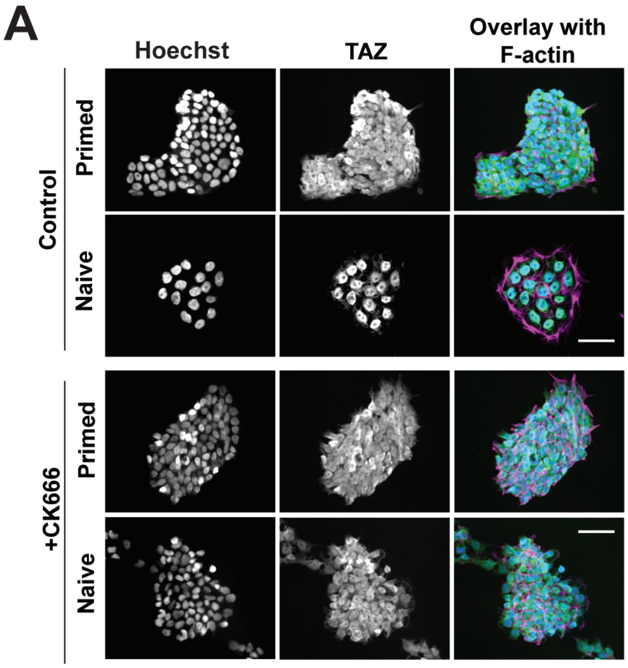


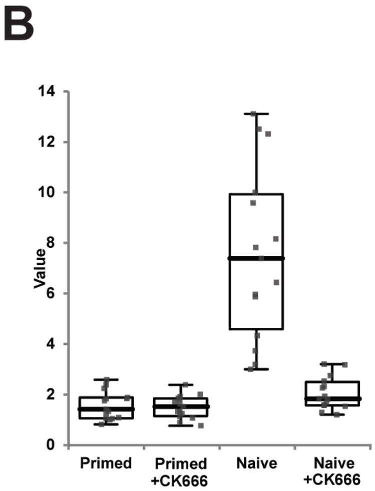
